# Passive Audio Vocal Capture and Measurement in the Evaluation of Selective Mutism

**DOI:** 10.1101/250308

**Authors:** Helen Y Xu, Jacob Stroud, Renee K Jozanovic, Jon Clucas, Jake Son, Bonhwang Koo, Juliet Schwarz, Arno Klein, Rachel Busman, Michael P Milham

## Abstract

Selective Mutism (SM) is an anxiety disorder often diagnosed in early childhood and characterized by persistent failure to speak in certain social situations but not others. Diagnosing SM and monitoring treatment response can be quite complex, due in part to changing definitions of and scarcity of research about the disorder. Subjective self-reports and parent/teacher interviews can complicate SM diagnosis and therapy, given that similar speech problems of etiologically heterogeneous origin can be attributed to SM. The present perspective discusses the potential for passive audio capture to help overcome psychiatry’s current lack of objective and quantifiable assessments in the context of SM. We present evidence from two pilot studies indicating the feasibility of using a digital wearable device to quantify child vocalization features affected by SM. We also highlight limitations in the design and implementation of this preliminary work that can help guide future efforts.

## AN OVERVIEW OF SELECTIVE MUTISM

Selective Mutism (SM) is an anxiety disorder characterized by persistent failure to speak in certain social situations but not others. SM is often diagnosed in early childhood when children are expected to start engaging in typical social interactions (Martinez et al., 2015). Children with SM are typically comfortable speaking in their home environment yet tend to struggle when challenged with novel social situations — particularly the school environment (American Psychiatric Association, 2013). SM-related symptoms can present long-term difficulties for children in developing social communication skills, in performing at school and in engaging with peers or others (Bergman et al., 2002). There is a growing need to objectively measure symptoms of SM to better understand the disorder.

## EVOLUTION OF THE SELECTIVE MUTISM DIAGNOSIS

Symptoms relating to SM were described as early as the late 1800s, when the disorder was referred to as “aphasia voluntaria” (Krysanski, 2003). The disorder was first captured in the psychiatric nosology of the Diagnostic Statistical Manual (DSM) in its third edition in 1980, wherein “elective mutism” was formally introduced. The criteria and interpretation of the disorder in DSM-III focused on a child’s *refusal* to speak, emphasizing beliefs that the disorder was rooted in defiance or trauma (American Psychiatric Association, 1980). The DSM-III-R removed the association between elective mutism and social phobia, recognizing them as separate conditions (American Psychiatric Association, 1987). As the field increasingly questioned the volitional nature of the disorder, the DSM-IV made two major changes: 1) the key diagnostic criterion was changed from a ‘*refusal* to speak’ to a ‘*failure*’, and 2) the name was changed to *selective* mutism. These changes deemphasized unwillingness and oppositionality and moved away from strictly psychosocial conceptualizations of SM; however, SM remained classified under “Other Disorders of Infancy, Childhood, and Adolescence” (American Psychiatric Association, 2000). Over the last decade, a growing consensus has pointed towards roots of SM in anxiety (Anstendig, 1999). In 2013, the DSM-V reclassified SM from childhood disorders to anxiety disorders (American Psychiatric Association, 2013).

## PREVALENCE

While SM was traditionally considered a relatively uncommon disorder, recent estimates suggest a prevalence of 0.47–1.0% of the population (American Psychiatric Association, 2013; Bergman et al., 2002; Viana et al., 2009/2); the increased estimates are thought to reflect a growing awareness of the disorder and prior misdiagnoses (e.g., autism, communication disorder, PTSD, or just “shyness”) (Lehman, 2002; Schwartz and Shipon-Blum, 2005). Although interest in SM is growing, research is relatively limited in comparison to research on other disorders of similar prevalence or severity (Bergman et al., 2013; Oerbeck et al., 2014). To date, much of SM research has consisted of case studies or intervention trials with small samples, making replication and generalization difficult.

## TREATMENT

SM has historically been considered difficult to treat, with symptoms often persisting long after treatment (Manassis and Tannock, 2008). SM treatments with published evaluation studies largely fall into four categories of treatment approaches: behavioral, psychodynamic, psychopharmacological, and systems-based (Zakszeski and DuPaul, 2017). Cognitive behavioral therapy (CBT) and selective serotonin reuptake inhibitors (SSRIs) have each demonstrated some efficacy and are commonly used; the effectivenesses of these treatments are consistent with conceptualizations of SM as an anxiety disorder (Carlson et al., 1999; Dummit et al., 1996). Research in the field of SM treatments has generally remained scarce, and more research is needed concerning different interventions (Manassis et al., 2016; Zakszeski and DuPaul, 2017). A focus on new and improved assessment methods in future research will help to illuminate why and how SM develops.

## THE CHALLENGES OF ASSESSMENT/CURRENT MEASUREMENT TOOLS

Diagnosing SM and monitoring treatment response can be quite complex. Currently no gold standard instruments or strategies exist for the quantification of SM symptoms. Diagnosis is largely dependent upon symptom reporting by parents, caregivers and teachers, often in the context of an unstructured diagnostic interview with a clinician. To assist in screening and diagnosis, some clinicians and researchers have introduced tools such as the ADIS, a semi-structured interview for anxiety disorders that has a Selective Mutism-dedicated module (Letamendi et al., 2008; Silverman and Albano, 2004). The Selective Mutism Questionnaire (SMQ) and the corresponding School Speech Questionnaire are commonly used to help quantify symptom severity (Bergman et al., 2001). Evaluations to rule out alternative diagnoses may include speech and language, oral-motor, and hearing assessments. Some providers carry out live behavioral observation sessions to gather data regarding how an affected child interacts with different individuals, including the parent (Child Mind Institute, 2016).

Currently, we lack objective, quantifiable assessments. Most assessments rely upon single raters and may be biased and limited in their application to SM. One approach in early exploration is surface electromyography (sEMG) for detecting changes in laryngeal tension, a suspected physiological response related to a failure to speak (Klein and Ruiz, 2017; Ruiz and Klein, 2013). Although potentially valuable, the applications of sEMG are limited due to the intrusiveness of placing electrodes on children’s necks. Sensory integration problems often cooccur with SM; therefore, methods that are less physically invasive and burdensome on the child are likely to be more effective (Schwartz et al., 2006). In addition, a need remains for other objective measurement tools of SM symptoms, such as the frequency and amplitude of vocalizations.

## CAN PASSIVE AUDIO CAPTURE PUSH THE FIELD FORWARD?

Passive audio/vocal capture is rapidly emerging as a promising assessment modality for psychiatry. In part, this trend reflects the availability of increasingly sophisticated analytic platforms for automated extraction of features, which can be used to predict states and behaviors (e.g., anxiety, depression, suicidality). Existing wearable technologies in consumer and research domains has already been successfully applied to monitor a range of behaviors and responses, including sleep, diet, electrodermal activity and heart rate (Crawford et al., 2015; Poh et al., 2010). The successes of devices such as the Fitbit and Apple Watch have helped to increase public acceptance of, and sometimes reliance on, wearable devices. Sensors for minimally intrusive audio capture have been employed in areas including stress research (Adams et al., 2014) and in nursing home monitoring (Rabbi et al., 2011). LENA, a device and software allowing for passive measurement of vocalization counts, vocal volume, and other conversational measures in children (LENA Research Foundation, 2016b), has been employed in recent speech studies, including studies of children with ASD and of bilingual children (Dykstra et al., 2013; Kashinath et al., 2015).

In a review of selective mutism research, Kratochwill recommends that “direct measures of speech [… and] physiological measures seem especially relevant in research and treatment of mutism. […] Such assessment may be especially relevant where medication is employed and/or where strong anxiety is prominent” (2014, 130–132). A benefit of passive, unintrusive devices for children with SM is maximizing their comfort, particularly because these children often become more anxious in new settings with new people. Simple passive audio tools could provide objective measures to better characterize the disorder without relying on complex analytics, burdensome devices, or multiple biased reports.

## A TEST OF FEASIBILITY: THE LENA DEVICE

Here we present findings from two initial tests of feasibility for the use of passive vocal recording to assess individuals with SM. We made use of LENA digital language processors (DLPs), the benefits of which include: 1) size smaller than a deck of cards, 2) availability of t-shirts designed to house a DLP in a chest pocket, 3) ready availability of automated feature extraction software and 4) ability to record and parse speech from the child and nearby speakers.

In both test applications, participants were provided a LENA shirt and DLP, which they decorated with name tags and stickers in an effort to acclimate the children to the shirts.

### Test 1: Brave Buddies

#### Overview

Brave Buddies is an intensive one-week SM treatment program at the Child Mind Institute that draws from a number of previously established behavioral techniques (i.e., adapted parent-child interaction therapy, group therapy and parent training). Our primary goals were to assess: 1) the ability of children with SM to tolerate wearing a LENA DLP for an extended period of time and 2) the ability of the DLP to detect relevant changes during the course of the intervention.

#### Methods

##### Participants

12 of 36 patients enrolled in Brave Buddies agreed to simultaneously participate in the LENA research study (9 female, 3 male; ages 5–8).

##### Design

Brave Buddies took place in a classroom-like setting and was structured like a typical school day, with each day divided into activity blocks (see Figure 1.IV). Patients were separated into three age-based groups of 12 children each, with one LENA study participant in the ages 4– 5 years group, five in the 5–6 years group and six in the 6–8 years group. Each group had its own room and dedicated counselors trained in behavioral techniques. Each child was also paired with a “Big Buddy” (counselor) who accompanied the child throughout each day’s activities and regularly prompted the child to answer questions and to vocalize. Throughout the treatment program, research staff accompanied each of the three groups, noting start and end times of each activity block, as well as information about deviations and factors that might affect the quality of the recordings.

**Figure 1.**
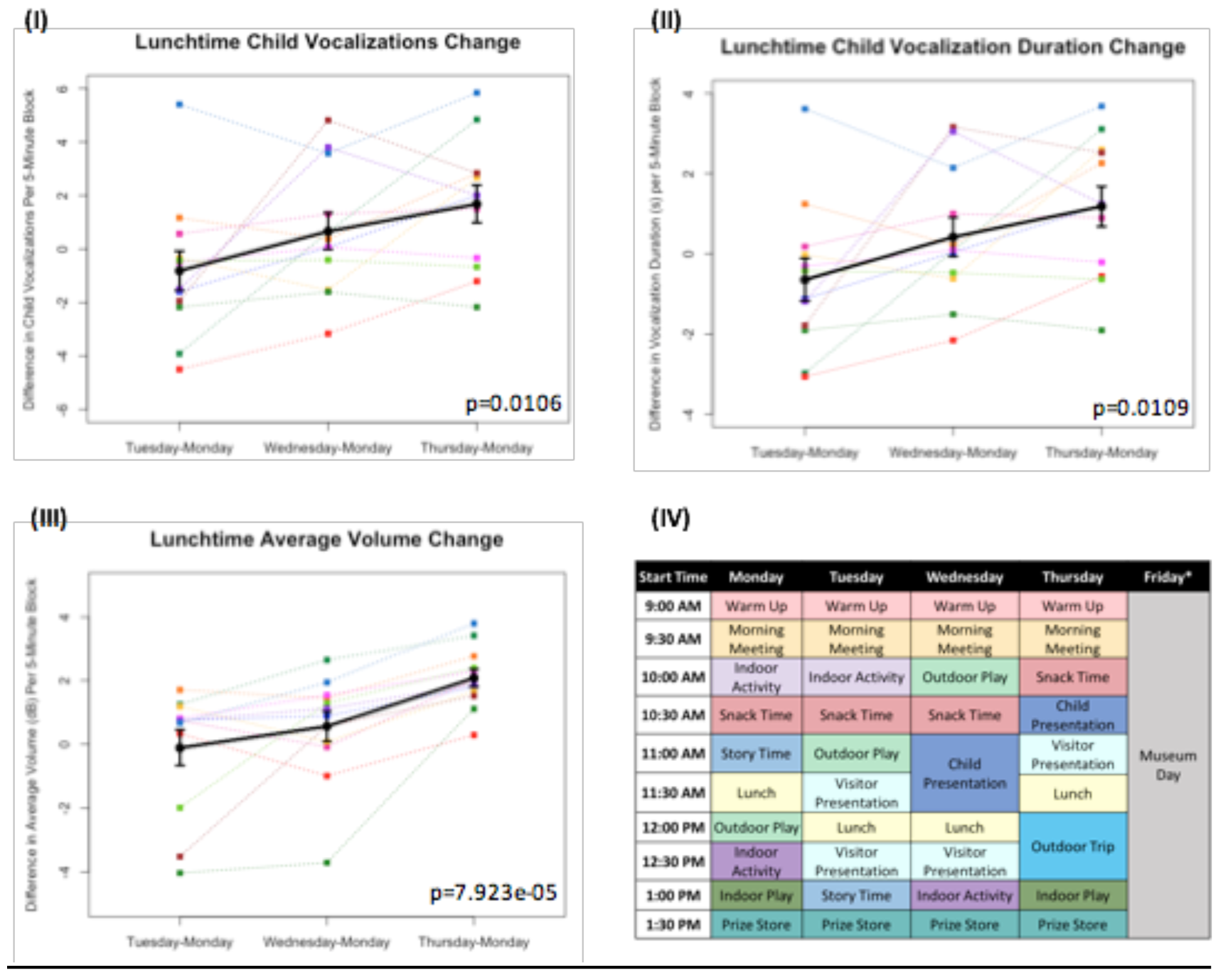
**(I), (II) and (III):** Differences in measures as compared to Monday’s baseline values plotted*. Each color line represents a different individual participant. Means with standard error bars plotted in black. (**IV):** Sample schedule for Brave Buddies week, showing various activities. *Friday data excluded from analyses, as described in Results.

##### Feature Extraction

LENA Pro, a software companion to the DLPs, provides numerous measures from the collected data (LENA Research Foundation, 2011; Xu et al., 2009); from these measures, we focused on a few measures of interest:

- *Vocalization counts:* instances of speech-related sounds separated by at least 300 ms of silence, produced by the child wearing the DLP.
- *Vocalization duration:* combined duration in seconds for all speech by the child per 5-minute block.
- *Average volume:* average decibel level per 5-minute block.
- *Conversational turns:* pairs of vocalizations between an adult and the child occurring within 5 seconds of each other. If either the child or adult responds to the other within 5 seconds, that is one turn.

#### Results

All 12 children were able to complete the five-day assessment of the LENA DLP without any significant difficulties related to wearing the device. Brave Buddies data from Friday, during which the children spent the day visiting a museum, were excluded due to a divergent setting and structure compared to previous days. From Monday–Thursday, we found that the most vocalizations occurred during the Outdoor Play activity as compared to other activities. In comparing data across days, we focused on Lunchtime, an activity that allowed for open, freeform vocalizing and was consistent across days. Using multivariate repeated measures ANOVA (scripts used available at https://github.com/ChildMindInstitute/LENA_BB_CPP_analysis/tree/master/BB), we found significant increases across the days in Lunchtime vocalization count (p=0.0106), vocalization duration (p=0.0109), and average volume (p=7.923e-05) per 5-minute block (see Figure 1.I–III); insignificant ANOVA results and no upward trends were observed in other activity blocks.

#### Discussion

Multiple detectable vocal properties exhibited significant improvements across the four days included in our examination (Monday–Thursday), though only during Lunchtime. The specific sensitivity of Lunchtime to changes in behavior may be informative; specifically, this was among the least structured and directed of activities, with less feedback and interaction from clinical staff. This finding suggests that there are limitations to simply applying a DLP to an ongoing intervention that does not specifically facilitate assessment of freeform speech. Looking forward, introduction of more such periods could increase the utility of passive audio capture in structured clinical intervention programs such as Brave Buddies.

There were two additional limitations of the LENA for tracking progress during Brave Buddies. First, despite its structure, the program involved numerous variables that were difficult to control from a research standpoint, such as lack of experimental controls, minimal free form speech, and lack of consistency in treatment applications. For example, the school-type activities in Brave Buddies were structured so that children would not be continually speaking, even if they participated a few times per activity, and treatment was based on individual needs and severity, conditions that vary across children. Second, participants in Brave Buddies were selectively biased towards less severe cases of SM who would be able to tolerate an unfamiliar group setting. Whether LENA and related wearables will be feasible with more severe populations remains unclear.

### Test 2: Controlled Play Paradigm

#### Overview

We carried out a more controlled assessment of the LENA DLP in which children were assessed one at a time and interactions more regulated. Specifically, we assessed children wearing a LENA while they were playing with their parent in an observation room in a design based on Parent-Child Interaction Therapy (Carpenter et al., 2014). Because a foreign environment alone may not be enough to evoke SM symptoms, we also varied whether a male experimenter was present and if present, whether he interacted with the child. The Controlled Play Paradigm was intended to test whether audio features extracted by the LENA software could differentiate children with SM from controls and to investigate correspondence between these features and established questionnaires (i.e., Selective Mutism Questionnaire (SMQ)).

#### Methods

##### Participants

12 children diagnosed with SM ages 5–8 (9 female, 3 male) participated, including 7 who also participated in Test 1 (Brave Buddies). 12 age-matched controls without any reported diagnoses were recruited from the community, ages 5–8 (7 female, 5 male).

##### Design

At the start of the timed study, the child and parent were left alone in a room filled with various toys (e.g., blocks, board games, toy animals, etc.). Research staff observed from another room via a one-way mirror. Speakers streamed audio into the staff observation room, and video was recorded with a view of the child and parent. After setup, video recording and LENA recording were started simultaneously, with both recording 5 blocks of exactly 10 minutes each. The parent or guardian also completed questionnaires, including the Selective Mutism Questionnaire (SMQ), which assesses child vocalizations in different settings, as well as interference and distress.

Three block types were included in an alternating block design (A-B-A-C-A). In Block A (no stranger), the parent was instructed to play with their child alone and to ask their child questions. Block B (stranger without interaction) introduced a male member of the research staff who had not yet interacted with the child as the “stranger.” He entered the room, told the parent and child, “I am going to do some work over here,” and sat in a corner of the room without further interaction. In Block C (stranger with interaction), the same “stranger” returned to the room, sat next to the parent and child and asked, “It looks like you’re having fun. Can I play with you?” The stranger engaged directly with the child, playing and asking questions (at least 2 per minute, often more). The parent was instructed to allow the stranger be the primary person asking questions during this block and to refrain from “saving” the child by answering questions intended for the child if the child failed to answer. During all blocks, the parent and stranger each wore an earpiece connected to a walkie talkie, through which observing research staff communicated.

#### Results

The data were divided by group (Control v. SM) and by condition (block A v. B v. C). Multivariate ANOVA showed no significant main effect of condition or interaction effect between group and condition for any measures. However, the main effect of group was significant for vocalizations (p=4.79e-07), vocalization duration (p=9.7e-06) and conversational turns (p=2.1e-07) (See Figure 2.I–III). A leave-one-out cross-validation of a generalized linear model predicting SM diagnosis from each of these measures resulted in an area of >0.7 under the receiver operating characteristic curve for each model, with most SMQ scores performing only slightly better (See Figure 2.IV–VII).

**Figure 2.**
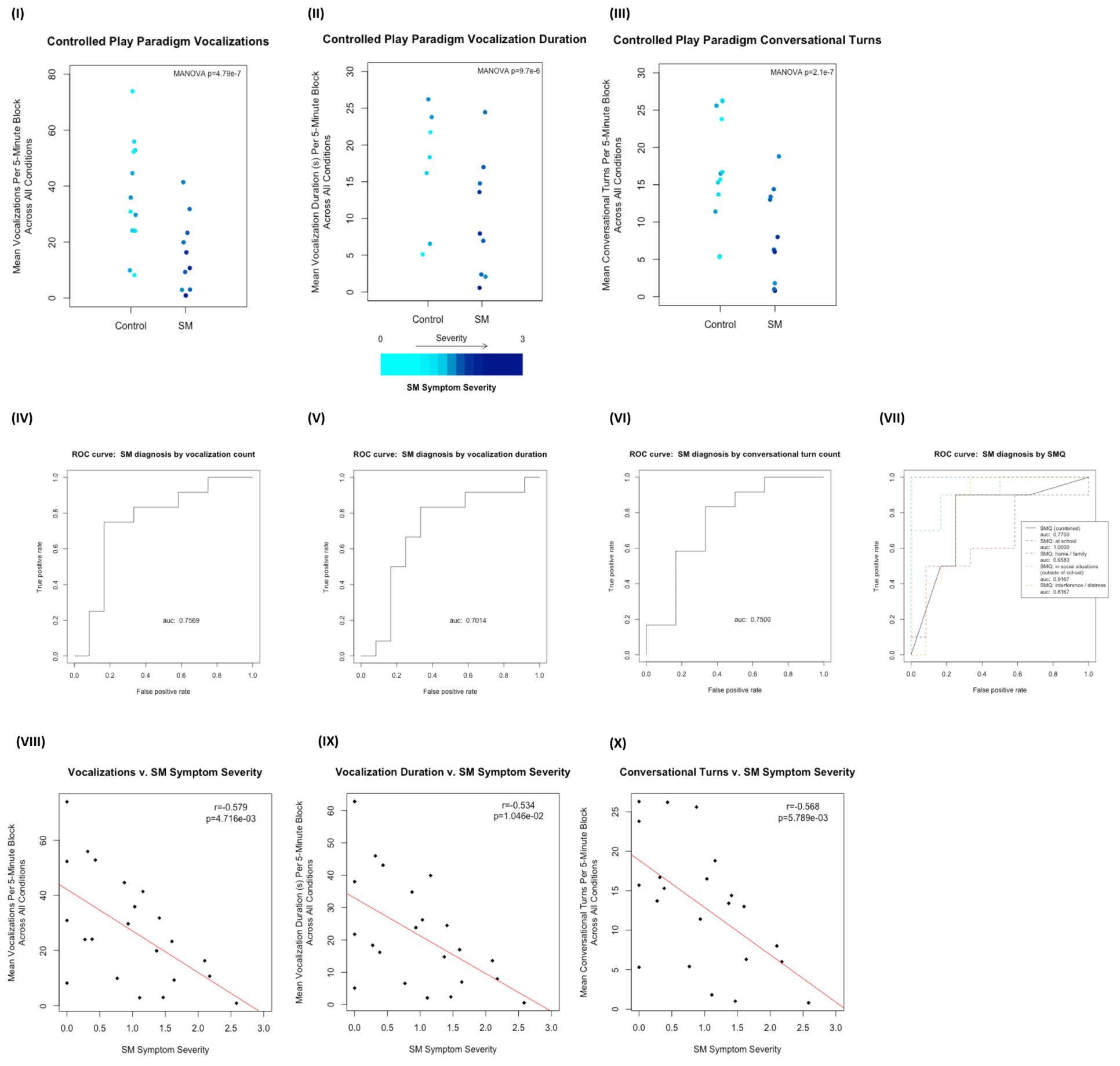
**(I), (II) and (III):** Control v. SM groups plotted with respect to mean vocalization counts, mean vocalization durations, and mean conversational turn counts across all conditions (A_1_, B, A_2_, C and A_3_ collapsed). Plotted points color scaled to the individual’s SM Symptom Severity score. **(IV), (V) and (VI):** ROC curves for leave-one-out cross-validation of generalized linear models predicting control v. SM group membership from each of the same measures. **(VII):** ROC curves for the same analysis of SMQ scores (combined and subscale) v. SM group membership. **(VIII), (IX) and (X):** Correlations plotted for same measures v. SM Symptom Severity for all 24 individuals. Line of best fit plotted in red.

SM Symptom Severity is a measure calculated based on SMQ responses from participants’ parents. SMQ Interference/Distress subscores (ranging from 0–18) were scaled and inverted to match the other subscores (ranging from 3–0) by this formula: *interference_score_scaled = 3 - (interference_score ÷ 6)*. The SM Symptom Severity was then calculated as 3 minus the mean score of the resulting 4 subscales (Home/Family, Social Situations, School and inverted Interference/Distress), representing an approximation of parent-reported SM-related symptom severity, with higher scores indicating increased severity. SM Symptom Severity was significantly negatively correlated with vocalizations (r=-0.579, p=4.716e-03), vocalization duration (r=-0.534, p=1.046e-02) and conversational turns (r=-0.568, p=5.789e-03) (see Figure 2.VIII–X).

#### Discussion

Within the controlled environment, we found the LENA DLP with LENA software could be used to detect between-group differences in various measures of vocalization. In each of the three scenarios, children with SM and control groups differed in mean vocalization counts per 5-minute block. A statistically significant linear relationship was demonstrated between SM Symptom Severity (calculated from SMQ responses) and each of three outcome measures extracted by the LENA software (i.e., vocalization count, duration and conversational turns). Thus, the LENA measures appears to be sensitive to SM-related changes in child vocalization, though further validation and refinement is needed before real-world clinical applications could be considered.

## CONCLUSIONS AND FUTURE DIRECTIONS

Selective mutism is an understudied anxiety disorder that would benefit from objective measures to characterize the heterogeneity of symptoms and treatment outcomes. This study indicates that the extraction of features from passive audio can be informative for SM research.

The LENA device is appealing for assessment of clinical populations, such as SM patients due to its availability and automatic processing; however, the device presents specific limitations for use with these populations. The LENA was developed for very young children, ages 0–4 years (LENA Research Foundation, 2016a), and though our work indicates its potential for older participants, those populations are not the developers’ focus. The LENA is also closed source and proprietary, meaning that its algorithms are unknown and immutable and we cannot know if our recordings are adequate for calibration. Lastly, the LENA is capable of recording *successful* vocalizations, but may not be able to detect *unsuccessful* or *extremely low-volume* vocalization *attempts*.

Moving forward, we will refine our experimental design based on lessons learned in this initial work, consider alternate or additional audio analysis options (Eyben et al., 2016; Sage Bionetworks, 2015) and develop more practical ways to use the LENA device for SM populations. The stimuli provided in each of these experiments did not provoke significant symptomatic behaviors from our participants; as such, future work may include more provocative stimuli (e.g., having a stranger offer a snack to probe for comorbid dysphagia).

As a behaviorally defined condition, SM appears to be derived from various heterogeneous factors (Hayden, 1980), and “given the complexity of the phenomenon labeled ‘selective mutism,’ it appears that multiple measures and their degree of correspondence are necessary” (Kratochwill, 2014, 132). Passive audio tools can provide multiple objective measures to better characterize SM and provide consistent feedback, empowering children and caregivers to better understand its etiology, to diagnose and to treat selective mutism in the future.

